# Feasibility of multimodal behavioral, gaze, and EEG measurement of natural scene segmentation in autism

**DOI:** 10.64898/2025.12.17.695033

**Authors:** Darnell K. Adrian Williams, Tridib K. Biswas, Concetta Brusco, Sophie Molholm, Ruben Coen-Cagli

## Abstract

Understanding how individuals organize complex natural scenes into perceptual segments is essential for explaining both typical and atypical sensory processing. In Autism Spectrum Disorder (ASD), differences in visual segmentation are widely documented. However, existing paradigms often rely on simplified stimuli and do not capture the dynamic, naturalistic processes underlying real world scene segmentation. To overcome those limitations, in this work we present a novel, multimodal experimental framework. The framework combines precisely timed, trial-based perceptual measurements with electroencephalography (EEG) to examine how neural activity aligns temporally with segmentation decisions, and with eye-tracking to examine visual exploration strategies and attention allocation. To demonstrate the framework, we report basic characterization results from a pilot study. Neurotypical (NT) participants and participants with ASD aged 16 years and above viewed natural scenes and textures and indicated whether two cued image regions belonged to the same or different perceptual segments. Each trial used a different pair of cues, and responses across trials enabled the estimation of subjectively perceived segmentation maps and the associated uncertainty. Both cohorts performed the task successfully and produced interpretable segmentation maps. Analysis of reaction time distributions, gaze profiles, and trial-aligned neural responses give us a window into cohort differences and the opportunity to study individual heterogeneity. Together, these findings establish a reproducible method for studying the temporal and spatial components of natural scene segmentation and illustrate that meaningful behavioral and neural distinctions in ASD may be studied with this approach. This framework provides a methodological bridge between controlled experimental stimuli and real-world perception in both neurotypical and neurodivergent populations.

## INTRODUCTION

Autism Spectrum Disorder (ASD)^✝^ is a highly diverse, heritable, lifelong neurodevelopmental condition defined clinically by two core diagnostic criteria: (1) persistent impairments in social communication and interaction, and (2) restricted repetitive patterns of behavior and/or interests^1,2^. Although not part of the diagnostic criteria, many individuals with ASD^✝^ also show numerous differences in sensory perception^1–6^, which are the focus of the present study.

These sensory features typically present throughout development, are prevalent in individuals with autism regardless of intellectual ability, and vary widely. These atypicalities include both hyper– and hypo-sensitivities to the world around them^6,7^ and include a range of symptoms, such as heightened sensitivity to sound or light and, in some cases, remarkable perceptual talents^8–10^. Individuals with autism are often less susceptible to visual illusions and frequently demonstrate exceptional and unique performance in visual search and discrimination tasks^4,6,11–16^.

These sensory and perceptual differences can meaningfully affect the daily life of people with autism. Theories such as the Weak Central Coherence^11^ and the Enhanced Perceptual Functioning^6^ propose that individuals with ASD tend to integrate visual information differently than neurotypical individuals. This refers both to integration of information across different parts of the visual field or between different visual features, as well as more broadly to integrating bottom-up, sensory-driven signals and top-down signals that convey prior expectations^17^. These classical theoretical frameworks have contributed to the interest in sensory symptoms, driven in part by the possibility that the core social features of autism may stem from more fundamental differences in sensation and perception^18,19^. As a result, there is strong motivation to understand the neurobiology and mechanisms underlying these sensory and perceptual atypicalities, ranging from interactions between brain areas^14,20–23^ to interactions between neurons in recurrent circuits^24–26^.

In this article, we adopt a recent experimental approach^27^ for studying visual integration, based on visual segmentation, which refers to the ability to distinguish objects from their background and from one another. ^28,29^ Segmentation is necessary to interpret complex visual scenes and thus fundamental to perception. It is shaped by both bottom-up sensory processing and top-down modulation^14,30–35^. Prior studies have shown that individuals with ASD exhibit atypical patterns of visual segmentation, including slower facial and texture segmentation and altered performance on embedded figures tasks^4,6,12–14,36,37^. These differences may contribute to broader alterations in visual perception and object-level processing. However, it remains unclear whether these effects extend to natural scenes, as most prior work has relied on highly simplified textures or coarse low-level stimuli. Examining segmentation in natural scenes is essential because it more accurately reflects the demands of everyday visual processing. This added complexity is thought to contribute to and shape some of the social and sensory differences often observed in ASD^18^. The first contribution of this article is to demonstrate that the experimental paradigm that was previously validated in neurotypical (NT) participants ^27^, enables a precise characterization of natural image segmentation in ASD.

Our second contribution is to adapt the experimental paradigm to include simultaneous measurements of eye movements and brain activity. Individuals with ASD frequently display irregular eye movements and altered fixation patterns for both social and non-social visual stimuli^38–42^. This suggests that differences in how visual information is sampled, prioritized, and integrated may relate to perceptual differences. Combining the gaze data collected in our paradigm with the segmentation maps will enable exploring this possibility and further characterizing visual saliency in ASD for both social and non-social stimuli. In addition, because perception and eye movements unfold rapidly and interactively, the ability to track neural activity in real-time is essential for understanding how sensory input is processed and integrated with attention and prior expectations^43–45^. This is particularly important in autism, where differences in sensory processing and perceptual integration are theorized to arise from altered coordination between sensory-driven and higher-order brain activity^46,47^. For this reason, the experimental design presented here also includes electroencephalography (EEG). EEG captures brain activity with millisecond precision, allowing us to examine the temporal dynamics of perception as they occur.

We employ a perceptual segmentation task^27^ in which, on each trial, participants are briefly presented with an image and they judge if a pair of cued locations on the image belongs to the same segment or different segments (Fig. 1). Data collected for several pairs of locations on one image enables the reconstruction of segmentation maps for each participant as well as the quantification of subjective uncertainty associated with the perceived segments (Fig. 2). We integrate these measurements with recordings of gaze behavior and EEG-recorded neural activity. Unlike prior work on segmentation in ASD, we apply this paradigm to natural images as well as to synthetic textures. This multimodal framework provides high-resolution temporal and spatial data to compare perceptual organization, attentional allocation, and brain-wide dynamics across individuals and stimuli.

**Figure 1.**
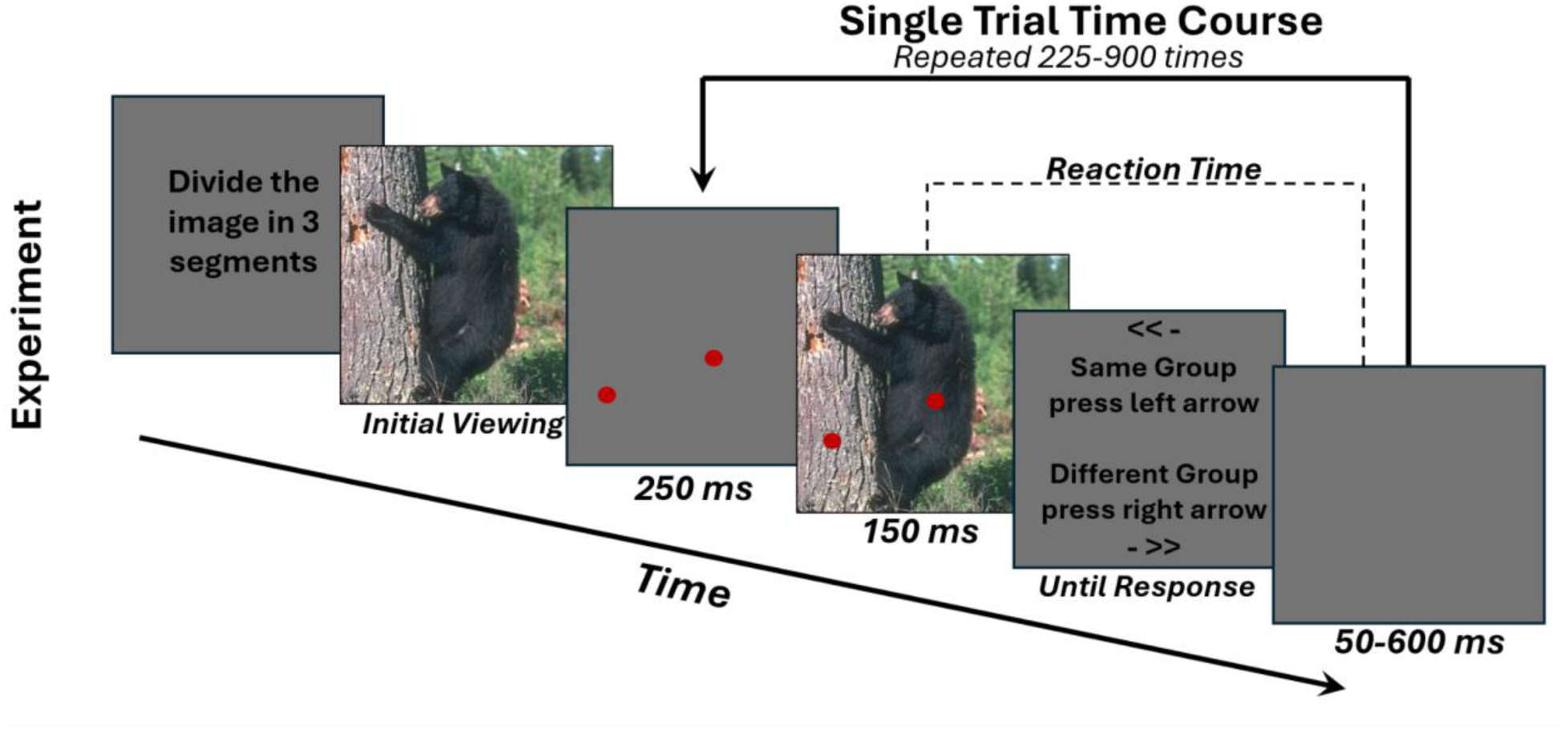
One experimental block of the segmentation task. Each block began with a text prompt instructing participants to mentally divide the scene into a predetermined number of segments. The image to be segmented was then displayed once and the participants viewed the scene freely. Next, a sequence of trials started. In each trial, two red circles were presented for 250 ms at pseudorandom locations on a grey screen, followed by the same image for 150 ms with the circles superimposed. Immediately afterward, a text prompt asked participants to indicate whether the parts of the image cued by the two circles belonged to the same segment or to different segments, using a left or right key press. Key mappings for “same” and “different” were randomized across participants but remained constant for each participant across trials and blocks. Reaction time (RT) was defined as the time elapsed between the offset of the image and the key press (participants were not allowed to respond before image offset). After the response, a grey screen was presented during an intertrial interval lasting between 50 and 600 ms.

**Figure 2:**
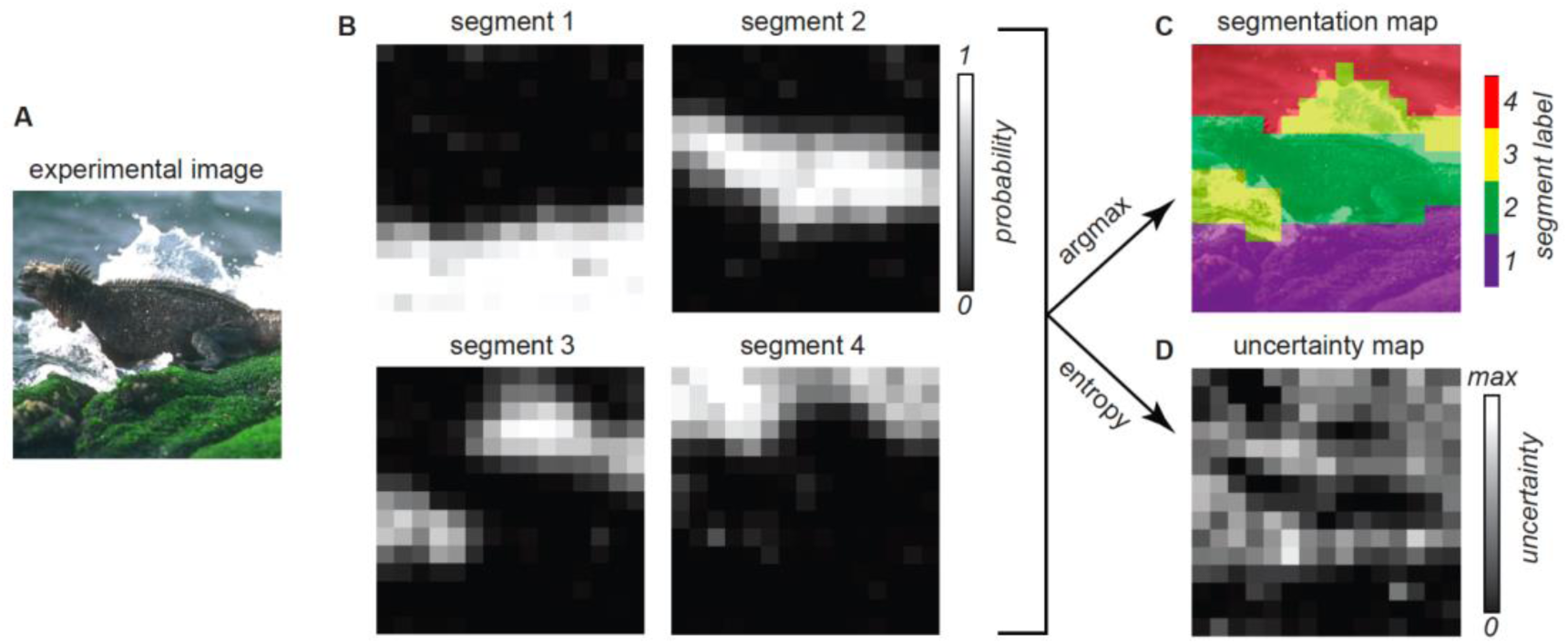
Example segmentation maps estimated from one experimental block. **(A)** The image used in this experimental block. **(B)** The probabilistic segmentation map estimated from the data of one participant. There is one map for each segment; within each map, the brightness of each pixel represents the probability that the pixel was assigned to the corresponding segment. **(C)** The segmentation map estimated by assigning each pixel to the segment with highest probability (i.e. using the argmax operator). Colors reflect segment identity, but not ordinal or semantic information. **(D)** The perceptual segmentation uncertainty map. Brightness corresponds to the entropy of the probability distribution over segment labels, which is a measure of perceptual uncertainty (brighter pixels denote higher uncertainty; the maximum possible value depends on the number of segments, K, and it is given by log(K)).

We demonstrate this paradigm with data collected in a pilot study with 24 NT and 13 ASD participants. We find that individuals with ASD are able to perform the segmentation task as well as NT participants, and their segmentation maps appear qualitatively comparable across both natural scenes and textures. We illustrate how our multimodal measurements can be used to test for differences in the timing and uncertainty of perceptual reports, visual exploration strategies revealed by spatial maps of eye movements, and the dynamics and spatial distribution of visually evoked neural activity. The resulting data, while preliminary, open the door to the possibility that differences in the perception of segments may not be entirely reflected in the segmentation maps but rather in the timing and uncertainty of percepts and in the underlying neural dynamics. These findings represent a useful entry point to investigate differences in perceptual segmentation in autism.

## METHODS

### Participants

We enrolled 24 neurotypical participants and 13 participants with an autism spectrum disorder diagnosis. The neurotypical group was approximately gender-balanced, with 11 of 24 participants identifying as female, whereas the ASD group was predominantly male, with only 2 of 13 participants identifying as female. Because ASD is diagnosed more frequently in males and may be underdiagnosed in females, this imbalance reflects a common challenge in ASD research. To preserve sample size, we retained the full cohorts rather than subsampling NT participants to match the ASD cohort’s gender distribution, noting this imbalance when interpreting cohort comparisons. The NT cohort ranged in age from 16.3 to 29.7 years (mean 22.0 土 4.9). The ASD cohort ranged in age from 16.4 to 31.1 years (mean 23.0 土 4.4). All participants provided online informed consent prior to participation. All participants were native English speakers with normal or corrected-to-normal vision. A clinical psychologist confirmed ASD diagnoses based on ADOS and developmental history.

The Institutional Review Board (IRB) of the Albert Einstein College of Medicine approved all study procedures. Participants were compensated according to institutional guidelines for their time in the laboratory. All procedures conformed to the ethical standards of the Declaration of Helsinki.

### Preregistration

This study was not preregistered.

### Behavioral data acquisition, preprocessing and analysis

#### Task description

Figure 1 illustrates schematically one block of the experiment (detailed in Vacher et al., 2023^27^) in which participants completed the task for one image. The structure of each block was as follows. At the start of the block, participants were presented with on-screen text explaining the task instructions, namely to segment the image into K perceptual segments (K ranging from two to five, across blocks), and to provide responses using the left and right arrow keys.

After the text with the instructions, the image to be segmented was displayed allowing participants to freely view the image and decide how to segment it. Next, a series of short trials started. In each trial, participants were shown a pair of red circles on a gray screen of average luminance for 250 ms, followed by the same image with the circles overlaid for 150 ms. Both the image and the circles then disappeared, and a response screen appeared, prompting participants to indicate whether the two dots belonged to the same segment or different segments using a left or right key press. The key mappings for “same” and “different” were randomized between participants but remained identical in all the blocks performed by each individual participant. The response screen remained visible until a response was made. The time elapsed between the offset of the image and the key press was recorded as the reaction time (participants were not allowed to respond before the offset of the image). After the response, a gray screen was presented during an inter-trial interval of random duration between 50 ms and 600 ms milliseconds, to minimize anticipatory responses (note that this interval includes also the time elapsed between key press and key release, which varied across trials and participants).

Before starting the experiment, the task was described verbally. Participants were told that there is no right or wrong segmentation, and that during the experiment they should base their answers on the segments they perceive. This was further illustrated by showing on a computer screen two example images and several possible segmentation maps of those images. Next, participants performed 25-50 practice trials to become familiar with the task structure. After confirming that they understood the task, participants performed up to 6 blocks in a single visit, with a unique image in each block (two participants from the ASD cohort completed two visits for a total of 12 blocks). Participants were offered short breaks every 3 minutes during a block and longer breaks between blocks. At the end of the experiment, participants were asked to describe verbally how they segmented the image. This information was recorded for each image.

The duration of a single block depended on the number of trials performed. This was set to the minimum number of trials needed to reconstruct a segmentation map with spatial resolution of 15×15 for the given number of segments K (using the equations derived in Vacher et al., 2023^27^). For K=2,3,4,5 participants performed 225, 450, 675, and 900 trials, respectively. The time to complete a block, including voluntary breaks, ranged from approximately 5 to 30 minutes.

#### Visual stimuli and presentation

Visual stimuli were presented on a monitor (Dell UltraSharp 1704FPT) that was gamma calibrated. The monitor was placed at a distance of either 67 cm or 96 cm (for sessions with EEG and gaze recordings) and the images were rescaled to cover an area of 8.7×8.7 degrees of visual field. The viewing distance was controlled by using a chin rest. Stimulus presentation was controlled with custom Matlab code that used the PsychToolbox 3 toolbox^48,49^.

Each participant completed up to four blocks with natural images and up to two blocks with synthetic, parametric textures in a single visit. Natural images were selected from the Berkeley Segmentation Dataset 500^50^ (BSD500), a dataset of 500 natural scene images used in computer vision segmentation research. Images were pre-screened based on human-drawn segmentations provided with the BSD500 dataset, to exclude images with more than 10 distinct segments. Next, images were selected manually, cropped to a size 256×256 pixels, and divided into four categories corresponding to the number of segments K requested at the start of the block. In this paper, we report data collected for up to five images from each category with up to five NT and up to five ASD participants per image.

For the blocks involving artificial texture stimuli, we generated the images by first partitioning the image frame into two or three regions that were randomly shaped but spatially compact. We then filled each region with a distinct but similarly oriented texture. Textures were bandpass Gaussian noise textures, created as described in Vacher et al^27^. Each texture was synthesized by convolving a white Gaussian noise image with a spatially dependent filter, which produced the desired spectral properties in each region and resulted in oriented textures with controlled differences in spatial frequency and orientation content.

In total, we used 18 unique images (12 natural scenes and 6 textures). Note that for some experimental blocks the perceptual and/or EEG and/or gaze data were excluded after preprocessing and quality control (detailed below), hence the number of unique images varies for different analyses and is reported in the corresponding text or figures.

#### Estimation of segmentation maps and perceptual uncertainty

The trial by trial responses collected during one experimental block were used to estimate the segmentation map of each participant and image. This was done using the method described in Vacher et al^27^. Briefly, the goal is to estimate the probability that each pixel in the image belongs to any segment. The number of segments is set to the same value that participants were instructed to use at the start of the block. Each segment is given an integer label, and the method uses numerical optimization to find the probabilistic assignments (termed probabilistic segmentation maps; Fig. 2B) that are most compatible with the responses of the participants. Next, the segmentation map is obtained by assigning each pixel to the segment label with the highest probability (Fig. 2C). Pixels with a probability close to 1 for one segment, and close to zero for all other segments, denote high perceptual certainty, i.e. the participant was highly certain in the assignment of that pixel. Conversely, pixels with a similar probability for all segment labels denote high perceptual uncertainty. This perceptual uncertainty was further summarized in the perceptual segmentation entropy map (Fig. 2D), which reports the entropy of the probability distribution over segment labels at each pixel. The estimation of the probabilistic maps was obtained using the Python implementation of software package Vseg^27^ with hyperparameter λ (which controls spatial smoothing of the map) set to 5.

#### Quantifying the similarity between maps

We defined an Information Gain (IG) measure of similarity between two segmentation maps, as follows. For each image, the mutual information between two segmentation maps (i.e. the maps from two distinct participants) was calculated:

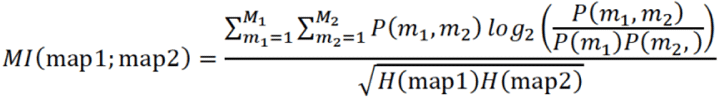

where *m*_1_ is the segment label in the first map (i.e. *m*_1_ ranges from 1 to the total number of segments *M*_1_in the first map) and similarly for *m*_2_; and *H* is the entropy of the map. The *MI* measures the similarity between maps because it quantifies how much the values of one map are predictable from the second map. This value was then compared to a null distribution of 10,000 values of *MI* between a randomly shuffled map (e.g. map 1) and the original version of the other map. This was done to construct a null distribution to determine significance of the *MI* value (one-tailed non-parametric permutation test, alpha=0.05). The shuffled maps were generated by randomly shuffling the spatial locations of map values, such that the marginal distribution of map values (*p*(*m*_1_) and *p*(*m*_2_)) remained the same. Therefore, we quantified the similarity between maps as the percent information gained from the map spatial structure relative to the null distribution. The same procedure was used to compute similarity between two perceptual uncertainty maps.

#### Exclusion criteria

In a small subset of experimental blocks (4/143 [2.8%] blocks for NT; 9/76 [11.8%] for ASD), the segmentation maps indicated that the participant had not performed the task correctly: data from those blocks was discarded. This was clearly indicated by segmentation maps that lack any spatial structure (e.g. Supp. Fig. 1A). Lack of spatial structure reflects that the responses were essentially random, i.e. unrelated to the position of the cues. We further confirmed this by estimating the segmentation maps from surrogate data where we randomly shuffled the measured responses relative to the cue pair presented in each trial: we observed that the segmentation map estimated from this surrogate data lacks spatial organization (e.g. Supp. Fig. 1B). Data from one additional block was excluded: after completing the block, the participant explained that they had partitioned the image into four equal quadrants rather than the segments they perceived (this was also evident from the estimated segmentation map; Supp. Fig. 1A).

#### Analyzing reaction times across blocks and participants

Before analyzing reaction times, we removed trials in which RTs were above the 90th percentile. We used median RTs as a measure of central tendency given that distributions were skewed. However, we found similar results using the arithmetic mean and the geometric mean. Confidence intervals around the median were found by bootstrapping with 9,999 resamples.

In Fig. 9, to compute the number of trials required for a participant to halve their RT (which we term the trial constant) we fit the parameters *α* and *τ* in the following equation:

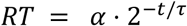

where RT is the reaction time, *α* is a scaling factor, *t* is the trial index, and *τ* is the positive-value trial constant. Parameters *α* and *τ* were fit using a nonlinear least-squares regression. To perform this analysis correctly, we needed to eliminate stimulus-specific or presentation-specific confounds. Therefore, we first z-scored the data: the mean RT for each unique experimental block (i.e. one participant seeing one stimulus) was set to 0, and the standard deviation to 1.

### Electroencephalography data acquisition and preprocessing

#### EEG Acquisition

For a subset of participants (11 in the NT cohort and 9 in the ASD cohort), we monitored neural activity with scalp EEG recordings. Participants were seated in a dark, electromagnetically shielded booth (Industrial Acoustics Company Inc., Bronx, NY). Neural activity was recorded using a 64-channel BioSemi ActiveTwo EEG system (BioSemi, Amsterdam, The Netherlands). The EEG signal was continuously sampled at 512 Hz, and analog TTL triggers marking the beginning of the trials and the participant responses were transmitted from the stimulus computer to the acquisition system and logged in a dedicated channel within the EEG recording (we term this the event channel). This information was used to define epochs and align EEG data temporally to task events and gaze data.

#### EEG Pre-processing and Bad Channel Handling

Raw data files in (.bdf) format were loaded into MNE-Python for pre-processing^51^. The event channel was preserved for event detection, and a standard BioSemi 64 montage was applied to assign 3D channel positions.

First, bad channels were detected using the PyPREP pipeline^52^, which employs a RANSAC-based detection algorithm to flag channels with high variance, flat signals, and poor correlations with neighboring channels. Visual inspection of channel variance plots was conducted for quality control. Identified bad channels were removed and subsequently interpolated from surrounding electrodes. Across files with available preprocessing metadata, the NT cohort had a mean of 2.00 bad channels per file (range = 0-3), while the ASD cohort had a mean of 2.25 bad channels per file (range = 0-4). A 60 Hz notch filter was applied to remove line noise, followed by a 1-120 Hz band-pass filter to attenuate low-frequency drift and high-frequency noise while preserving ERP-relevant activity. This range was chosen as a conservative threshold, as scalp EEG potentials rarely exceed amplitudes of +/− 100 μV as do the associated frequencies of potential interest^53^.

#### Event detection and epoching

Epochs were created from −500 ms to 1000 ms relative to trial onset. This extended window was intentionally chosen to fully encompass both the participant’s response period and the variable inter-stimulus interval that preceded the next trial, ensuring that as much task-relevant activity as possible was captured in each epoch. Subsequent analysis was restricted to a narrower window, to remove activity that may be related to the response in the preceding trial or to the onset of the subsequent trial. No baseline correction was applied at this stage to preserve raw signal characteristics for subsequent preprocessing.

#### ICA Artifact Removal

To remove artifacts from eye movement, muscle activity, and other non-neuronal sources, independent component analysis (ICA) was performed using FAST ICA with unlimited components, random state 42, and max 1000 iterations^54–56^. ICA components were visually inspected using the time courses of the components. Artificial components identified by spatial topography consistent with blinks, muscle activity, or channel artifacts were manually selected for exclusion by the experimenter. Across files with ICA metadata available, the NT cohort had an average of 1.50 removed ICA components per dataset (range: 0–4), while the ASD cohort had an average of 1.75 components removed (range: 0–5).

#### Epoch rejection and data retention

Following ICA correction, automated epoch rejection was performed using Autoreject^57^. Autoreject uses cross-validation to learn channel-specific noise thresholds and automatically repair or reject contaminated epochs to minimize human bias and improve processing quality. The algorithm tested for one, two, or four channel interpolations per epoch and selected the optimal strategy to maximize data retention while maintaining signal quality. Overall, AutoReject rejected on average 9.45% of epochs due to data quality issues (NT: 6.84%, ASD: 12.81%). Because the rejection rate was well below 50% in all experimental blocks, no blocks were excluded due to these flags. Clean epochs were saved in MNE Python (.FIF) files for subsequent analysis, and pre-processing metadata were saved as JSON files for tracking and quality control.

In addition to signal quality, a subset of epochs with unusually long RT were excluded from subsequent analysis because they could denote lapses in attention or periods when the participants took a voluntary break without communicating it. Specifically, epochs corresponding to the slowest 10 percent of RTs were excluded, followed by exclusion of all epochs with RTs exceeding 2.5 seconds.

#### Event Related Potential (ERP) Generation

ERP preprocessing was performed separately for each experimental block, i.e. for each participant and image. Neural activity in each epoch and channel was re-referenced to the global average (subtracting the mean of all EEG channels at each time point), then averaged across epochs. Baseline correction was applied to each channel by subtracting the mean amplitude of that channel within a pre-stimulus window extending from –100 ms to 0 ms. Because in some trials the inter-trial interval could be less than 100 ms, the estimate of the baseline activity could be contaminated by activity related to the previous trial’s response. To ensure that this did not affect the data presented in Results, we repeated all analyses using a shorter window of –25ms to 0.0ms which does not include the previous trial response, and obtained similar results.

For the analyses of Section 2.2, we averaged the ERP across selected channels to create a single ERP per image and per participant. In other analyses, we further aggregated data across different participants that segmented the same image, resulting in one ERP per image. Lastly, in other analyses we aggregate across different participants and images, resulting in a single ERP. The aggregation was always conducted separately for NT and for ASD participants.

#### Topographic Maps

To study the topography of ERPs, topographic maps were generated from individual evoked responses using all available EEG channels. Grand-average topographic maps were computed separately for NT and ASD cohorts using MNE-Python’s grand-average function. We also computed NT – ASD difference maps. All maps were displayed on a consistent –10 to +10 µV color scale, baseline-corrected from –100 to 0 ms, referenced to a global average reference, and filtered using the reaction-time exclusions explained above.

#### Latency analysis

For each participant and stimulus class, ROI peak latency was defined as the time of the maximum absolute ERP across O1, Oz, O2, PO7, PO3, POz, PO4, and PO8 within 0–500 ms after cue onset. GFP peak latency was the time of maximum GFP in the same window. Cohorts were compared with two-sided Welch tests. These latency tests were distinct from cluster-based permutation tests of amplitude over time; cluster boundaries were not interpreted as response-onset estimates.

### Gaze data acquisition and preprocessing

#### Eye tracking

For a subset of participants (16 subjects, for a total of 82 blocks in the NT cohort; and 4 subjects for a total of 26 blocks in the ASD cohort), we monitored gaze using an EyeLink 1000 system (SR Research, Mississauga, ON, Canada) at a sampling frequency of 500Hz. A 9-point calibration was performed at the start of every experimental block. Gaze data were collected throughout the entire block and analyzed separately in different epochs as explained below.

#### Preprocessing

Gaze data were preprocessed and analyzed using MNE-python. During preprocessing, blinks and gaze data outside of the image area were removed. Sessions were manually checked for quality: those with poor calibration and/or gaze density that appeared unrelated to the image were excluded (18/82 blocks for NT, 13/26 blocks for ASD).

Each experimental block comprises different periods, for which gaze data was collected and can be analyzed separately. These include: reading the text instructions; initial viewing of the image; trials epochs (between cues onset and response, see Fig. 1). Analysis reported in this paper focuses on gaze density during the initial viewing of the image. Gaze density was calculated from the gaze samples’ spatial coordinates using Gaussian kernel density estimation over the image area with a smoothing standard deviation of 10 pixels. Gaze density maps were then downsampled to the same spatial resolution as the segmentation maps (i.e. 15×15).

### Statistical significance

Different types of statistical significance tests were required for different data modalities and analyses. Therefore, the details are provided in Results.

## RESULTS

### 1. Spatial characteristics

#### 1.1 Perceptual segmentation maps

Our first goal was to determine if participants in both cohorts completed the task successfully. For each participant and image (termed a block), we used the numerical procedure described in Methods to reconstruct the perceptual segmentation map (Fig 2C) from the response data collected in the corresponding experimental block. Figure 3 demonstrates qualitatively that meaningful maps could be reconstructed for participants in both cohorts. The maps show alignment with recognizable objects in the natural image (e.g. the lizard, rocks, and water in Fig. 3, top) and with boundaries defined by local orientation differences in the texture (Fig. 3, bottom). As a control, we confirmed that no meaningful maps could be reconstructed when participants’ responses were randomly reshuffled across trials to make them unrelated to the visual cues (Supplementary Fig. 1B). Across our dataset, meaningful maps could be reconstructed for both cohorts in the vast majority of the experimental blocks (139/143 blocks for NT; 67/76 for ASD). In the remaining blocks, map reconstructions revealed that the participants responded randomly, i.e. they ignored the task instructions (Supplementary Fig. 1A).

**Figure 3.**
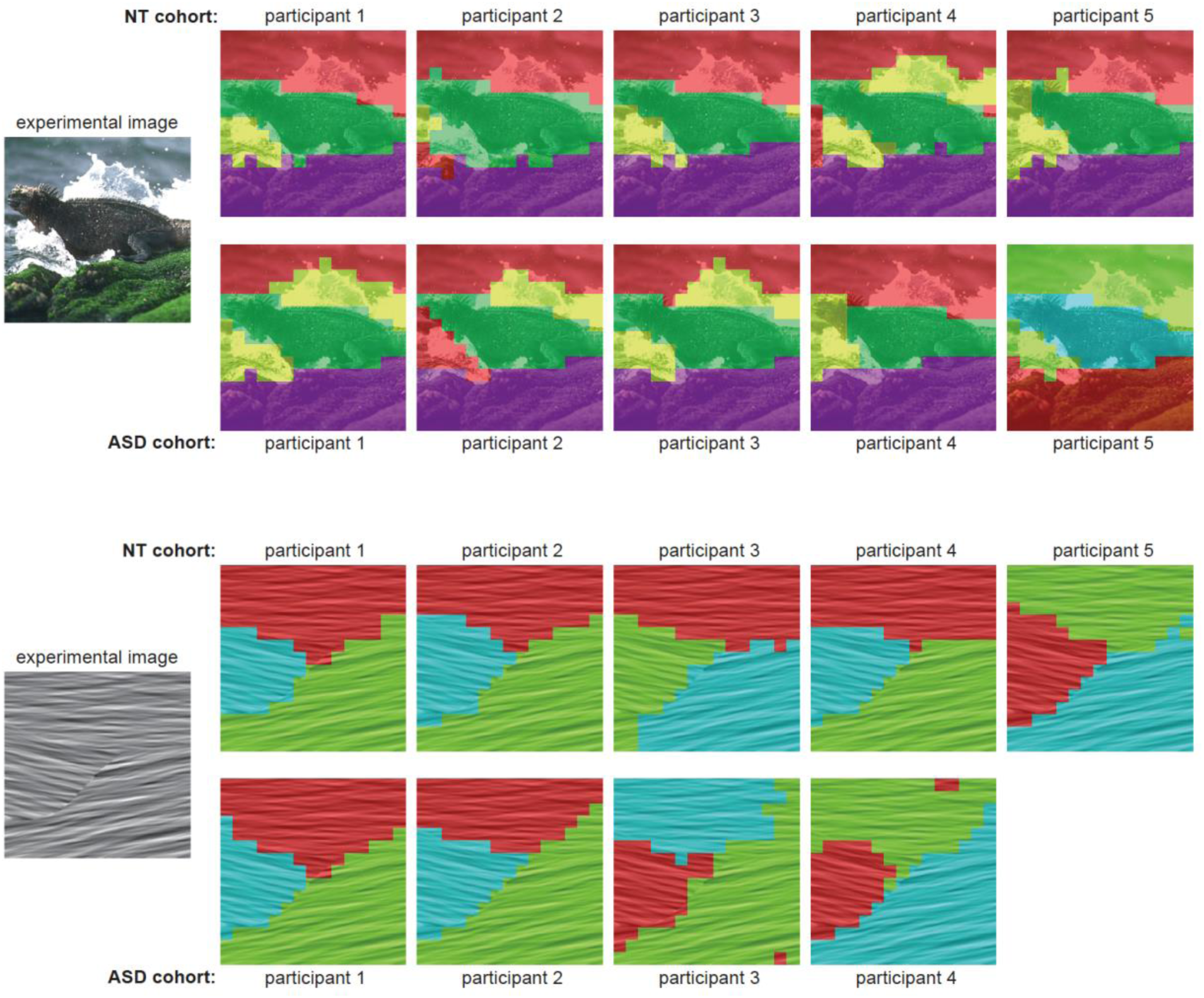
Example segmentation maps. Top, a comparison of perceptual segmentation maps for 5 NT participants and 5 ASD participants segmenting the same natural image (shown on the left). The colors corresponding to segment labels are overlaid on the image to facilitate visualization. The numerical values of the labels, and corresponding colors, do not convey ordinal or semantic information and thus should not be compared across participants. **Bottom,** A comparison of segmentation maps for 5 NT participants and 4 ASD participants segmenting the same synthetic texture image (shown on the left).

Although the maps were similar between individuals and cohorts, there were also noticeable differences in the details. For instance, in Fig. 3, top, some participants treated the water splash as a single segment on its own, others merged it with the background water, and others yet split the splash into two separate segments due to visual occlusion. To quantify the similarity of the segmentation maps between participants, we defined a measure termed Information Gain (IG; detailed in Methods): given two segmentation maps, the IG denotes how well the spatial organization of one map can be predicted from the other. A large percent IG value indicates that two maps are more similar between themselves than to surrogate maps in which pixel locations are shuffled randomly. An IG of 0% means the two maps are not more similar than two random maps.

We compared quantitatively between cohorts the inter-participant similarity of the maps using IG. Figure 4A illustrates one image for which the IG values were larger between pairs of NT participants than between pairs of ASD participants (mean IG = 14,383 +/-821% s.e.m. between all pairs of maps from 5 NT participants; mean IG = 6,494 +/-1,915% s.e.m. between all pairs of maps from 5 ASD participants). Across the entire dataset, we observed large values of IG (>100%) in both cohorts, denoting that the maps were highly consistent between participants (Fig. 4B). For natural images, the inter-participant similarity was marginally lower for ASD than NT (mean IG 5871+/-492 across 97 pairs for ASD, 7329+/-581% across 108 pairs for NT, t-test p=0.06; Fig. 4B). In contrast, for textures there was no difference between cohorts (mean IG 12561+/-1467 across 31 pairs for ASD, 12745+/-1165% across 46 pairs for NT, t-test p=0.92; Fig. 4B).

**Figure 4.**
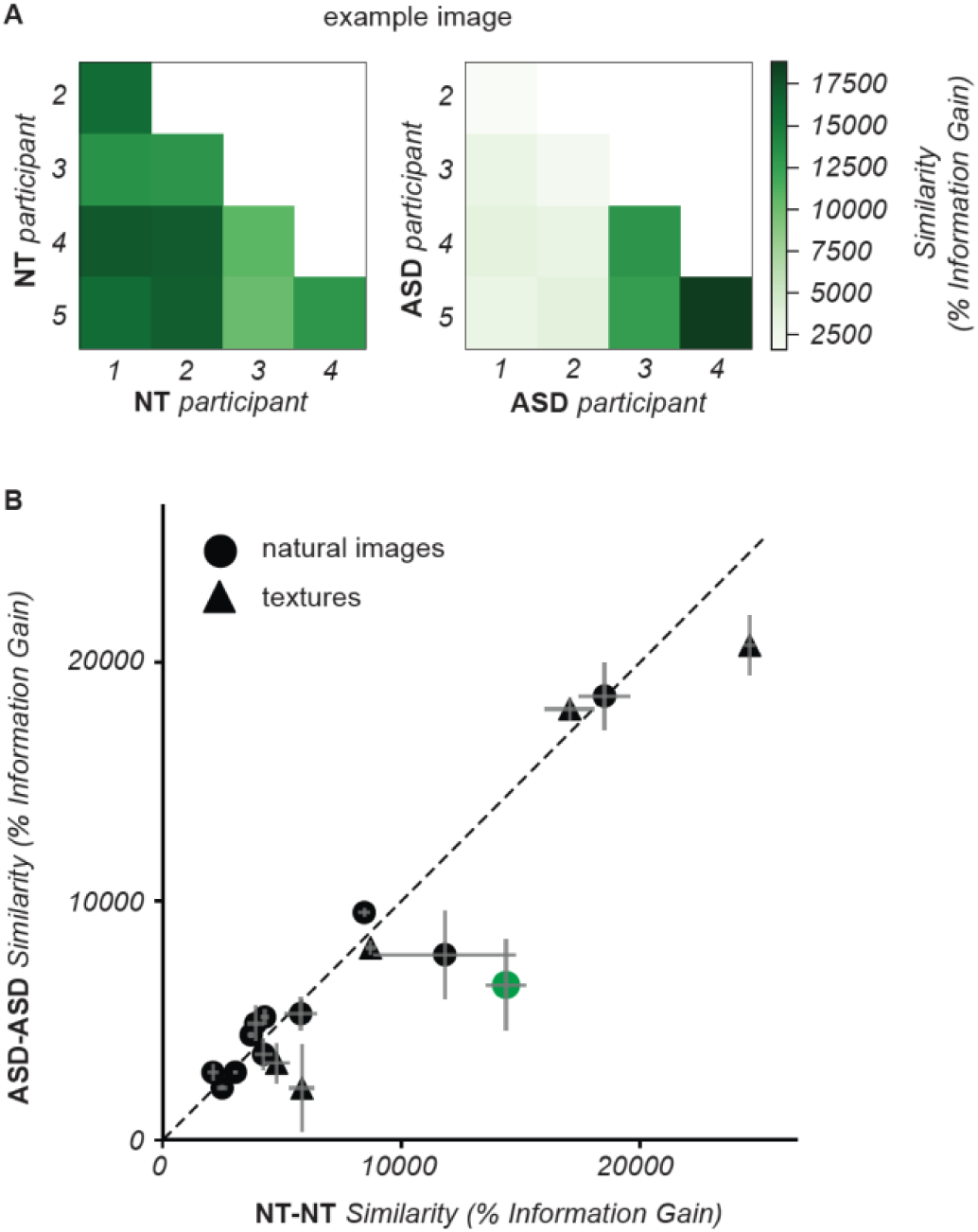
Similarity between the segmentation maps of different participants. **(A)** Each entry in the matrix represents the similarity between the segmentation maps of a single image by two participants in the NT cohort (left) and in the ASD cohort (right). Similarity was measured by information gain (IG), namely how much better the segment labels in one map can be predicted from the other map than from randomly shuffled maps (see methods). Darker shades of green correspond to higher IG, i.e. higher similarity. **(B)** Each symbol represents the mean IG for one image, averaged across all pairs of NT (abscissa) and all ASD (ordinate) participants that segmented that image. The green symbol corresponds to the example image in **(A)**. Error bars indicate the standard error of the mean across pairs of participants per image.

In summary, qualitative inspection of the segmentation maps indicates that both cohorts performed the segmentation task successfully and perceived meaningful segments in the images. Quantitatively, there was high similarity in the spatial organization of perceptual segments between individuals. For natural images, but not textures, there was a tendency for lower similarity between ASD than NT participants.

#### 1.2 Perceptual uncertainty

The segmentation maps analyzed in the previous section represent the most likely subjective interpretation of the image. Our measurements also convey richer information about the perception of segments. The reconstruction procedure (detailed in Methods) computes a probabilistic segmentation map (Fig. 2, left) from the trial-by-trial responses: namely, for each pixel we obtain an estimate of the probability that that pixel was assigned to any one of the possible segments. A pixel with a probability of 1 for a given segment, and 0 for all other segments, indicates that the participant was highly certain about that pixel. On the other hand, a pixel with equal probability for multiple segments denotes high uncertainty. To quantify this subjective uncertainty, we calculated the entropy of the probability values per pixel (higher values of entropy correspond to higher uncertainty; detailed in Methods). Figure 5 illustrates the segmentation uncertainty maps for the same images and participants as Fig. 3. Just like the segmentation maps, the uncertainty maps were qualitatively consistent between individuals for both the natural image and the texture.

**Figure 5:**
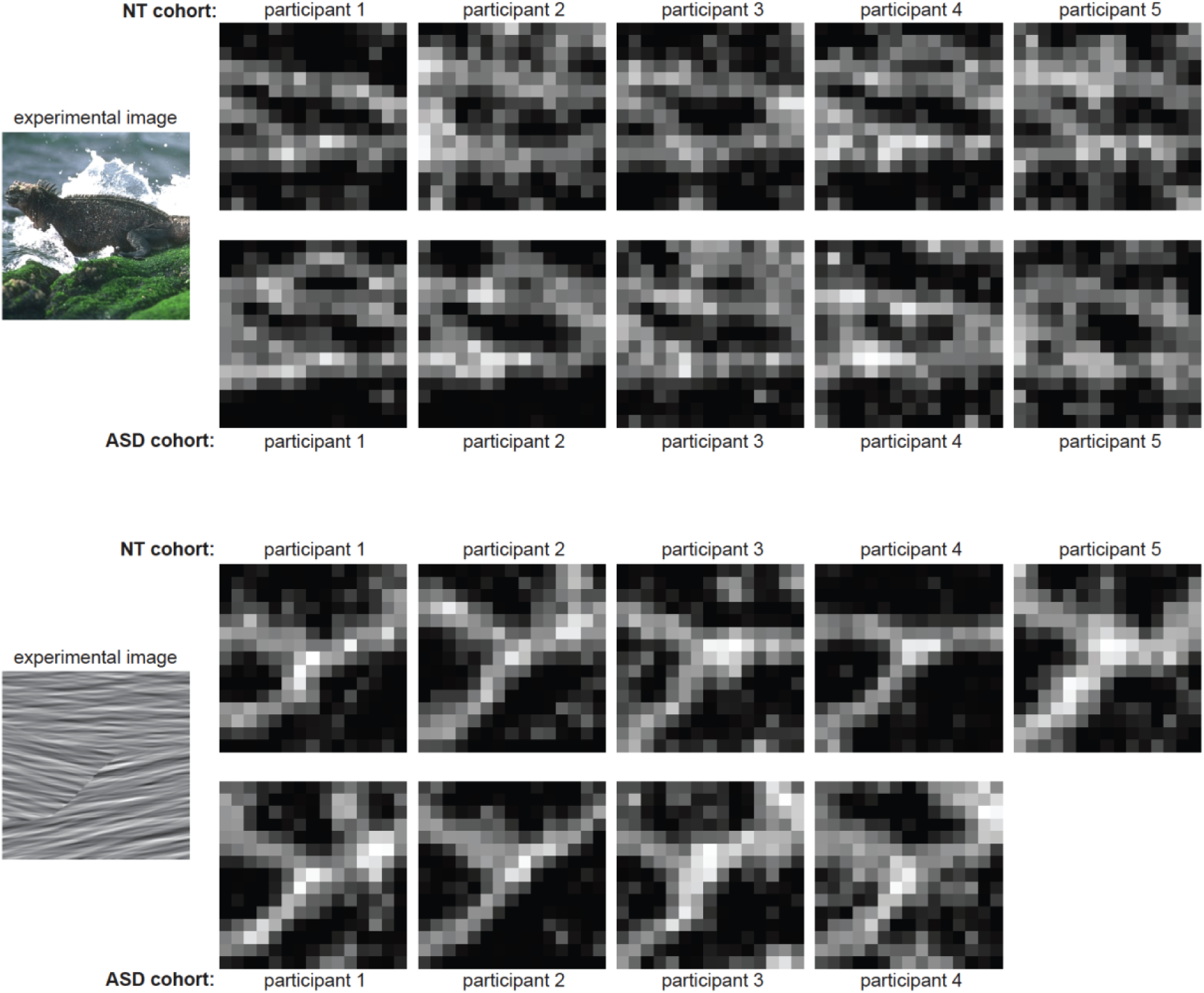
Example uncertainty maps. Uncertainty maps (defined in Fig. 2D) are shown for the same images and participants of Fig. 3.

We summarized each uncertainty map into a single value by computing the average across all pixels (normalized to the maximum possible uncertainty for the given number of segments). This quantity represents the overall subjective uncertainty for that particular image. For natural images, the overall uncertainty level was higher for ASD than NT participants (mean uncertainty 0.66+/-0.01 across 61 cases for ASD, 0.63+/-0.01 across 91 cases for NT, t-test p=0.012; Fig. 6A). Conversely, texture images induced lower uncertainty in the ASD cohort, although the difference between cohorts did not reach significance (mean uncertainty 0.67+/− 0.02 across 26 cases for ASD, 0.69+/-0.01 across 42 cases for NT, t-test p=0.21; Fig. 6A).

**Figure 6.**
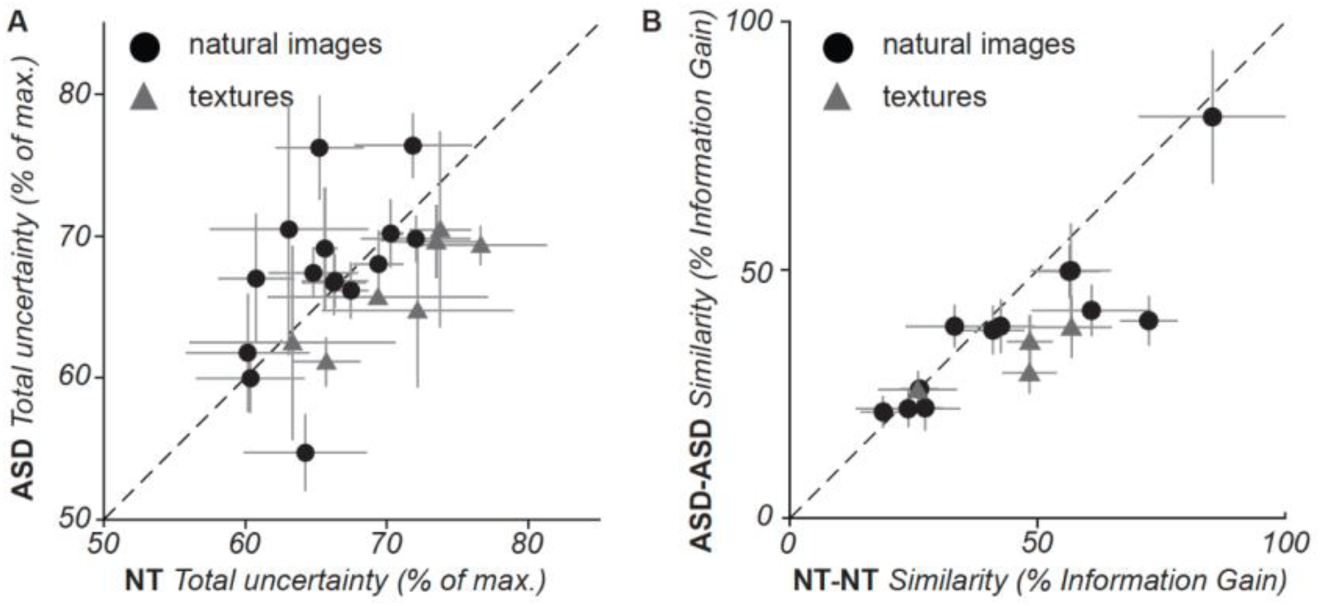
Comparisons of perceptual uncertainty between participants and cohorts. **(A)** Each symbol represents, for one image, the total uncertainty across all pixels in the image averaged across all NT (abscissa) and all ASD (ordinate) participants that segmented that image. Error bars indicate the standard error of the mean across participants per image. **(B)** Each symbol represents the mean IG between uncertainty maps for one image, averaged across all pairs of NT (abscissa) and all ASD (ordinate) participants that segmented that image. Error bars indicate the standard error of the mean across pairs of participants per image.

We then compared the spatial organization of uncertainty, using the same IG metric as for the perceptual segmentation maps. We observed that for the majority of images the spatial distribution of uncertainty was more similar between NT participants than between ASD participants (Fig. 6B). The difference was significant for textures (mean IG 34+/-3% across 22 cases for ASD, 47+/-3% across 31 cases for NT, t-test p=0.005) but not for natural images (mean IG 41+/-3% across 64 cases for ASD, 48+/-3% across 86 cases for NT, t-test p=0.13).

In summary, our measurements of uncertainty provide additional dimensions to characterize the subjective nature of perceptual segmentation and suggest differences in how NT and ASD participants segment natural images versus textures.

#### 1.3 Gaze maps, spatial coverage/spread

Eye movements are frequent perceptual decisions that vary person-to-person, moment-to-moment, and are highly driven by image content/salience. To study how these perceptual decisions may vary between cohorts, we recorded eye-tracking data in a subset of experimental blocks (82 blocks in the NT cohort and 26 blocks in the ASD cohort). Each block begins with an initial viewing period of the image to be segmented (Fig. 1, left); the amount of time spent in this epoch is determined by the participant. On average, ASD participants viewed the image for 10.58 +/-7.2 seconds (st. dev.); NTs viewed for 8.96 +/-5.18 sec.

To characterize how visual attention was allocated to different parts of each image, we computed gaze density maps which measure view time per pixel. Figure 7 demonstrates example gaze densities during the initial image viewing epochs of two participants (rows) viewing 5 images (3 natural and 2 texture images; columns). For instance, when viewing the image of a face (Fig. 7 column 1), the NT participant’s gaze was concentrated around the face’s eyes, whereas the ASD participant viewed other face regions. This appears consistent with known differences in ASD viewing behavior in social contexts^39^. Which image regions were the most salient perceptually (i.e. highest gaze density) also differed between the two example participants in some non-social natural contexts (e.g. landscape image, Fig. 7 column 2) and texture images (e.g. Fig. 7 column 4), but was qualitatively similar in others (Fig. 7 columns 3,5). These examples illustrate how this paradigm can be used to explore the possibility that differences in ASD visual saliency processes extend beyond social contexts.

**Figure 7:**
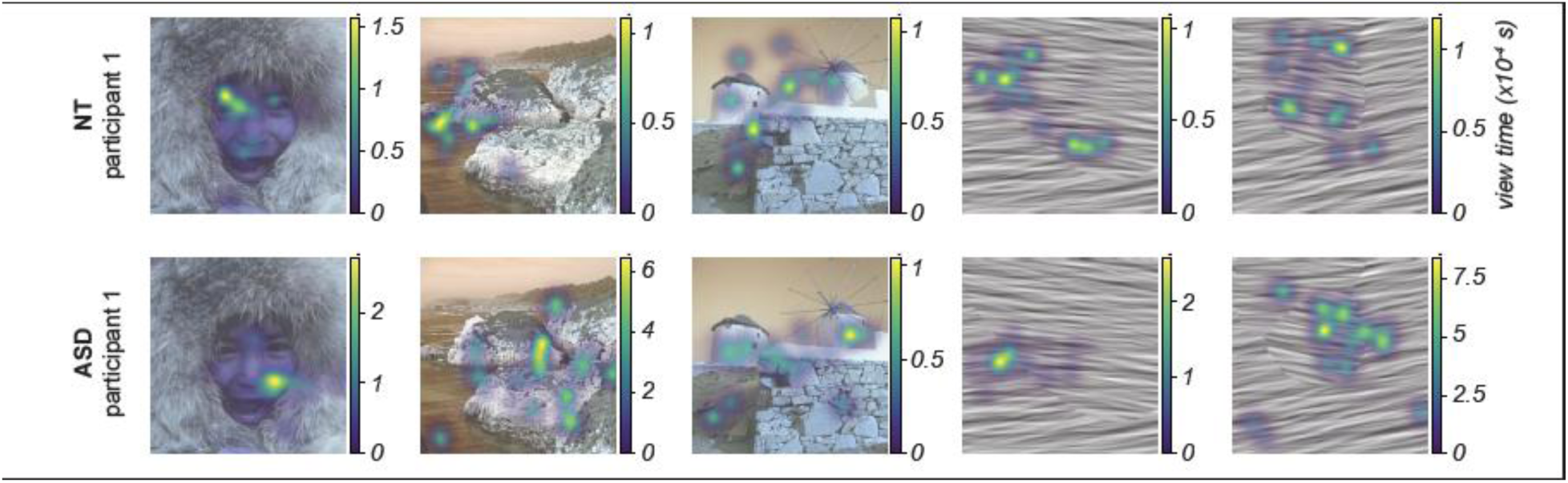
Example gaze density maps. Gaze density is represented by the colored overlay on each image and was computed as the total amount of time that gaze spent on each pixel of the image, during the initial viewing of the image for up to 10 seconds (Fig. 1, left). Each map was calculated using Gaussian kernel density estimation over all image pixels and a smoothing factor (sigma=10 pixels).

To quantify the spatial allocation of visual attention regardless of image content, we measured what fraction of spatial area of each image was viewed by each participant (% image coverage). Across all images, we found that ASD participants tended to view a greater spatial area of the image: Gaze densities during the entire initial viewing epoch on average covered 38.5+/-3.0% of the image for ASD participants (n=26 blocks), vs 31.5 +/-1.4% for NT participants (n=82 blocks) (two-sided t-test p=0.038). We asked if this difference could be explained by differences in viewing epoch duration, or if greater spatial coverage in ASD is also present on smaller time scales. To address this, we calculated gaze density and % image coverage at increasing time periods in the epoch (Fig. 8A), and found that the cohort difference was present as early as the first second of image viewing (Fig 8B). Thus, our results suggest differences in ASD spatiotemporal gaze dynamics on short, moment-to-moment timescales as well as during longer periods of image viewing.

**Figure 8:**
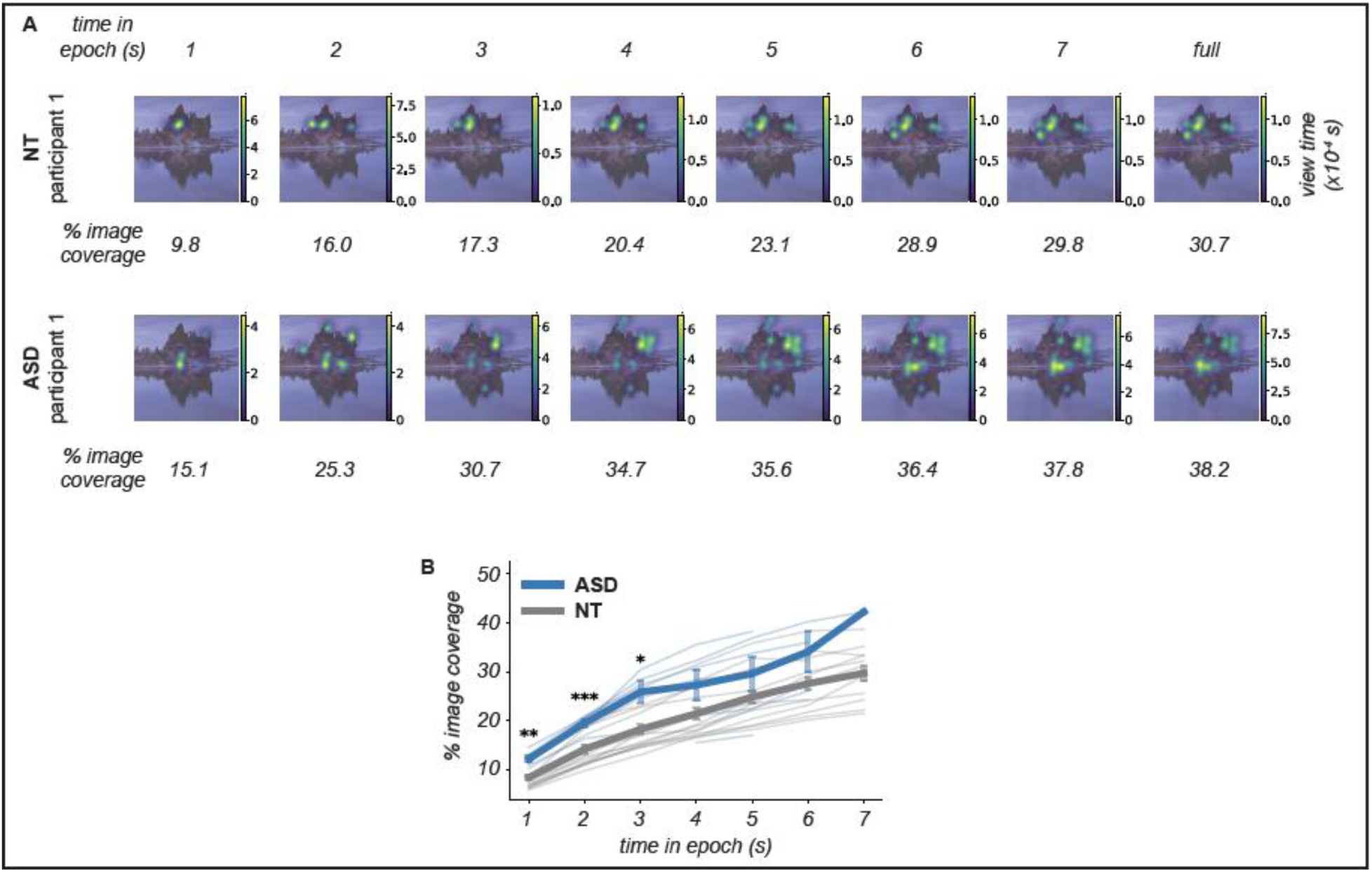
Comparison of image coverage by gaze during initial image viewing. **(A)** Gaze density is represented as in Fig. 7, for one example image. Each panel uses gaze data from a cumulative time window beginning at time 0 and extending for a duration indicated in the top row (time in epoch). The numbers underneath each panel quantify the percentage of the image area that has been visited by the gaze during the corresponding time window. **(B)** Each thin line represents, for one image, the mean across all participants in one cohort that viewed that image. Thick lines represent within-cohort means. Error bars indicate the standard error of the mean across participants and images, within each cohort. Note that total view time was determined freely by each participant, therefore the mean is taken over a smaller number of participants at longer times in epoch. *p<0.05; **p<0.01; ***p<0.001.

**Figure 9.**
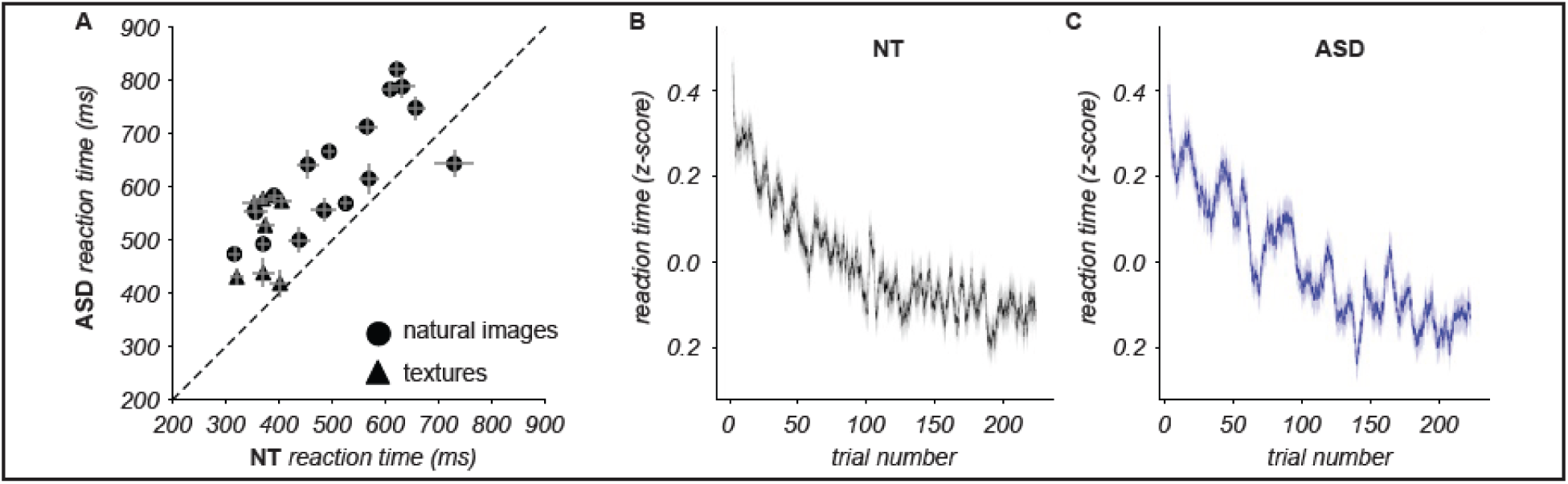
Comparisons of reaction times. **(A)** Each symbol represents, for one image, the mean RT averaged across trials in the experimental block and all NT (abscissa) and all ASD (ordinate) participants that segmented that image. Error bars indicate the standard error of the mean across participants per image. **(B)** RT for the first 225 trials across all experimental blocks for NT (left) and ASD (right) participants. RTs were z-scored per experimental block before combining across blocks. Trials were ordered by their trial number within each block (i.e. there were many trials indexed “trial 1” across all stimuli). A sliding window of 500 was used across all of these trials. Mean: The solid line and shaded area indicate, respectively, the mean and standard error of the mean within the sliding window.

### 2. Temporal characteristics

#### 2.1 Reaction times

A common finding in ASD research on perceptual decision making is that ASD participants often respond more slowly than NT. Interestingly, the opposite pattern is sometimes observed in tasks related to segmentation such as the Embedded Figures task^58^, the Wechsler Block Design^14,59^, and visual search of a target among distractors (a form of figure-ground segmentation)^60,61^. In our task, the RTs were compatible with the more commonly reported effect: Across all trials the median RT was 582 ms (bootstrapped std. err. 19.4 ms) for the ASD cohort versus 491 ms (bootstrapped std. err. of 43 ms) for the NT cohort (Fig. 9A). The difference between cohorts was significant both for natural images (582+/-43 ms for ASD, 491+/-19 ms for NT, Mann Whitney test p<0.005) and textures (468+/-28 ms for ASD, 334+/-32 ms for NT, Mann Whitney test p<0.0005). Although the absolute RT values varied by a factor of two or more within each cohort, the difference between cohorts did not appear to scale with the RT, i.e. the difference did not increase for images that took longer to segment.

One potential concern when comparing the RTs between cohorts is that ASD participants could have greater difficulty maintaining focus on the task throughout the entire duration of an experimental block. If that was the case, their RTs should increase as the block progresses, or perhaps oscillate as attention wanes and wades. To rule this out, we compared between cohorts how RTs changed over trials throughout the block. In both cohorts, RTs decreased monotonically with trial number (Fig. 9B). This finding indicates that there was no decline in focus or engagement throughout the block; on the contrary, participants rapidly improved their performance. To compare this effect quantitatively, we fit an exponential function and computed a “trial constant” enumerating, in each cohort, how many trials participants required to halve their RT (details in Methods). The trial constant for the ASD cohort was shorter for natural images (19.38+/-0.03 for ASD, 19.92+/-0.02 for NT, Mann-Whitney test p<0.0001) but longer for textures (21.2+/-0.03 for ASD, 17.1+/-0.02 for NT, Mann-Whitney test p<0.0001). However, an entire block lasted between 225 and 900 trials, therefore the difference between cohorts was less than 1% of the total number of trials in the block, indicating that both cohorts learned the sensorimotor contingencies on a similar time scale.

In summary, our measurements of RTs reveal a systematic and sizable difference in the dynamics of perceptual segmentation between cohorts. This difference cannot be attributed to a disengagement from the task or deficit of attention in the ASD cohort, indicating that our task reveals meaningful differences in perceptual processing and decision making.

#### 2.2 Neural dynamics in visual areas

In a subset of participants (n = 11 NT, 9 ASD) we recorded EEG data while they were performing the task. Because segmentation is known to involve recurrent interactions within occipital and parieto-occipital regions^14,62,63^, we examined activity from occipital and parieto-occipital electrodes (O1, Oz, O2, PO7, PO3, POz, PO4, PO8). We focused on event-related potentials (ERPs) in those electrodes by aligning activity to trial onset. Details of ERP calculation are provided in Methods. Given the brief cue–image sequence (400 ms), few saccades are expected, although fixation was not enforced. Their potential contribution to ERP morphology remains a consideration (see Discussion).

Figure 10 shows the ERP waveforms for both cohorts (11 NT participants, 64 blocks; 9 ASD participants, 57 blocks; see Supplementary Fig. 2 for single-image ERPs). Both cohorts exhibited a prominent positive-going deflection that peaked approximately 300–450 ms after cue onset. The cues appeared on a gray screen from 0 to 250 ms and were then superimposed on the image until 400 ms. Compared with ASD, NT participants showed a larger-amplitude response over the post-cue interval for both natural scenes and textures (Fig. 10A,B).

**Figure 10.**
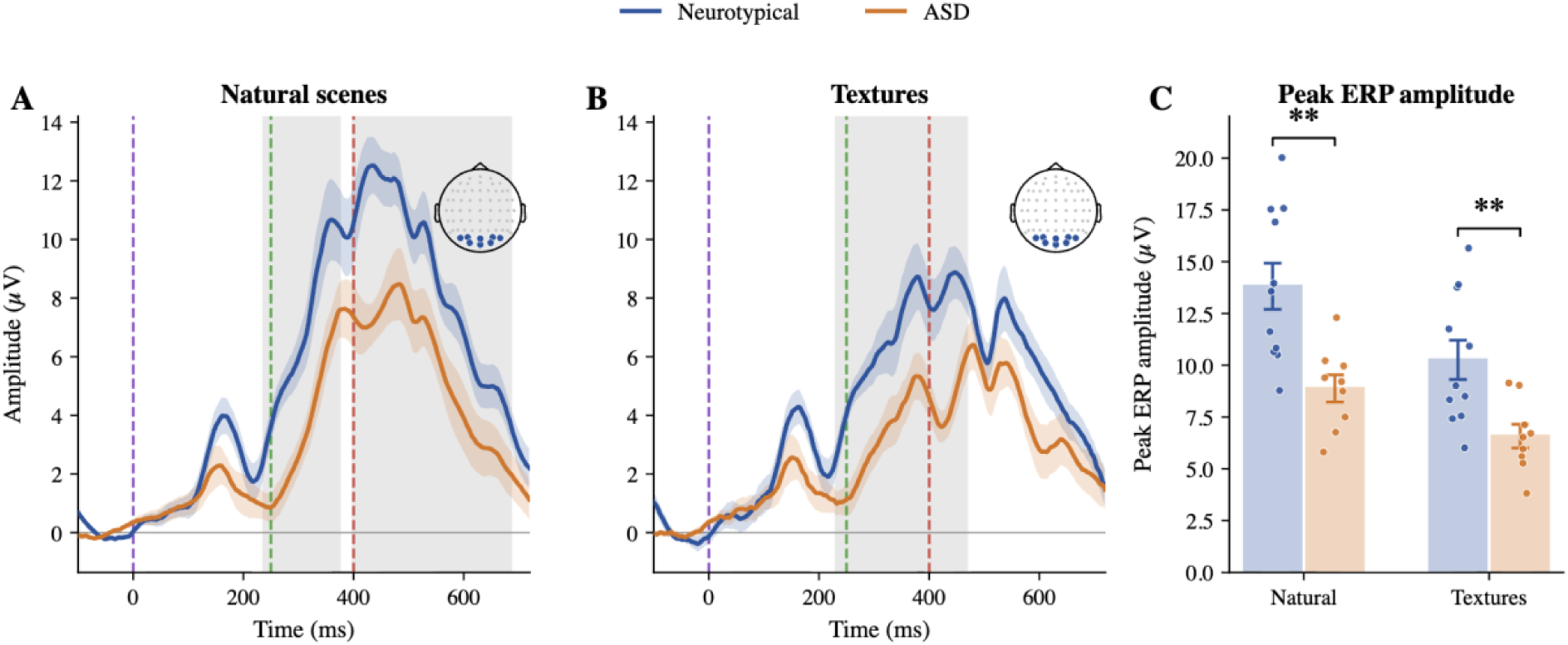
Grand-average event-related potentials over occipital and parieto-occipital sites. ERPs from O1, Oz, O2, PO7, PO3, POz, PO4, PO8 (highlighted on the inset head) for natural scenes (A) and textures (B), averaged across participants within each cohort (Neurotypical, blue; ASD, orange). Shaded regions denote the standard error of the mean across participants. Dashed vertical lines mark cue onset (0 ms, purple), image onset (250 ms, green), and image offset (400 ms, red). Gray bands indicate significant cohort clusters (cluster-based permutation, 4096 permutations, cluster-forming threshold F = 2.5, corrected across time): natural scenes 234–377 ms (p = 0.039; and a later 396–688 ms cluster, p = 0.021) and textures 229–471 ms (p = 0.007).

To evaluate the full time course while correcting for multiple comparisons across time, we used cluster-based permutation testing (independent samples, 4096 permutations, cluster-forming threshold F = 2.5) on subject-level ERP waveforms, separately for natural scenes and textures. ERP amplitude was higher in NT across significant clusters for natural scenes (234– 377 ms, cluster p = 0.039; 396–688 ms, cluster p = 0.021) and textures (229–471 ms, cluster p = 0.007). These corrected time-course tests, rather than separate tests at a selected peak, provide the primary inference for the ERP amplitude difference.

To evaluate response timing independently of amplitude-cluster boundaries, we defined each participant’s ROI peak latency as the time of the maximum absolute occipital/parieto-occipital ERP within 0–500 ms and compared cohorts with two-sided Welch tests. We did not detect a cohort difference for natural scenes (NT 423.7 ± 15.8 ms; ASD 444.4 ± 19.1 ms; p = 0.413) or textures (NT 404.3 ± 26.6 ms; ASD 432.7 ± 18.4 ms; p = 0.392).

In summary, the NT cohort showed larger-amplitude ERPs over occipital and parieto-occipital channels during and after image presentation. We did not detect a cohort difference in independently estimated ROI peak latency.

#### 2.3 Brainwide neural dynamics

To complement the prespecified occipital/parieto-occipital ERP analysis, we next characterized response magnitude and scalp distribution without restricting the analysis to that electrode set. These analyses asked whether cohort differences extended across the scalp and whether their distribution remained posterior.

To statistically quantify cohort differences in the spatial pattern of response amplitude across the scalp, we computed Global Field Power (GFP) continuously at every time sample (as for the ERP analysis). GFP reflects the spatial standard deviation of the scalp electric field at each time point and provides a channel-agnostic index of the overall strength and spatial differentiation of the neural responses^62^. NT–ASD differences were assessed separately for natural scenes and textures using cluster-based permutation tests across time (4096 permutations; cluster-forming threshold F = 2.5). GFP was higher in NT across significant clusters for natural scenes (221–719 ms, cluster p = 0.006) and textures (211–471 ms, cluster p = 0.012). Peak GFP within 0–500 ms was also higher in NT than ASD for natural scenes (8.33 ± 0.53 vs 6.14 ± 0.57 µV, p = 0.012) and textures (6.22 ± 0.49 vs 4.37 ± 0.42 µV, p = 0.010). For the latency analysis, each participant’s peak latency was defined as the time of maximum GFP within 0–500 ms. We did not detect a cohort difference in GFP peak latency for natural scenes (NT 432.5 ± 15.1 ms; ASD 443.4 ± 17.3 ms; Welch p = 0.644) or textures (NT 415.7 ± 12.3 ms; ASD 433.8 ± 19.9 ms; Welch p = 0.451). Thus, amplitude, not detected peak timing, distinguished cohorts in this dataset. Because the GFP difference preceded the delayed keypress, it is unlikely to reflect motor execution; without a task-matched control, however, we cannot attribute it specifically to segmentation rather than general visual processing.

To characterize where the cohorts differed across the scalp, we generated topographic maps for each cohort and NT – ASD difference maps (details in Methods). Figure 12 shows the NT–ASD difference maps at the latencies indicated above each scalp map. At each sampled latency, we tested the NT – ASD topography with a cluster-based permutation test across electrodes using the F statistic, correcting for multiple comparisons in space. Significant posterior clusters were present at 262 ms (cluster p = 0.002) and 327 ms (cluster p = 0.011); the 262-ms result remained significant after Bonferroni correction across sampled latencies. These effects were concentrated over occipital and parieto-occipital electrodes. These spatial tests identified electrode clusters at individual latencies but did not estimate their temporal duration. At 720 ms the difference map suggested a frontal pattern, but the corresponding spatial test was not significant; we therefore treat this later pattern as descriptive.

Together, these analyses show a spatially cohesive cohort difference over posterior occipital and parieto-occipital regions during visual processing. The temporal GFP test identified when overall field strength differed, whereas the electrode-cluster test localized the scalp distribution of the difference at 262 and 327 ms. This convergence supports a cohort difference in response magnitude and posterior scalp distribution.

## DISCUSSION

We have presented a novel paradigm to study perceptual segmentation of natural images in autism. Segmentation of visual inputs is a fundamental operation of the visual system, enabling observers to organize complex scenes into meaningful parts. Despite its importance, this process has traditionally been studied using simplified, synthetic stimuli, which limits our ability to understand how segmentation unfolds in realistic environments. Natural scenes more closely reflect the complexity of everyday visual input. Studying segmentation under these conditions provides an opportunity to understand how individuals with ASD organize complex visual information into meaningful parts.

We demonstrated that participants in both cohorts understood and completed the segmentation task, and that the paradigm can be combined with concurrent eye tracking and EEG. The pilot data identified cohort differences in reaction time, gaze coverage, ERP and GFP amplitude, and posterior scalp topography, together with stimulus-dependent differences in uncertainty organization. In contrast, we did not detect cohort differences in ROI or GFP peak latency. Given the small sample, these results establish feasibility and generate testable hypotheses rather than definitive evidence for an autism-specific neural mechanism. By combining trial-level psychophysics with natural and controlled stimuli and multimodal recording, the paradigm supports both exploratory and hypothesis-driven studies of perceptual organization.

Our paradigm builds on a design used in prior studies of perceptual segmentation, namely the “same segment / different segment” task^27,68,70,89^. Different from those past studies and other work on segmentation in autism, the paradigm we adopted enables measuring the spatial organization of perceptual segments and their associated uncertainty, through estimation of the segmentation maps (Fig. 2). One clear demonstration of the importance of obtaining these new measurements, is our finding that although perceptual uncertainty (Fig. 6A) appears similar between NT and ASD cohorts, the corresponding spatial maps vary across participants and reveal more idiosyncratic patterns of uncertainty in ASD participants (Fig. 6B). Given our parallel finding that ASD participants explore the images with more spatially distributed eye movements (Fig. 8), it will also be important to study how the spatial organization of segments, uncertainty, and gaze relate to each other and to neural variability^71^.

ASD participants responded more slowly than NT participants for both natural scenes and textures (Fig. 9A), consistent with past studies of decision making in ASD.^4,14,58^ The EEG analyses likewise identified cohort differences in response magnitude and posterior scalp distribution: NT participants had higher ERP amplitude in corrected temporal clusters (Fig. 10), higher GFP across corrected clusters and at the 0–500-ms peak (Fig. 11), and posterior spatial clusters at 262 and 327 ms (Fig. 12). Direct ROI and GFP peak-latency tests did not detect cohort differences. Together, these findings highlight cohort differences in response magnitude and posterior scalp distribution in this pilot sample.

**Figure 11.**
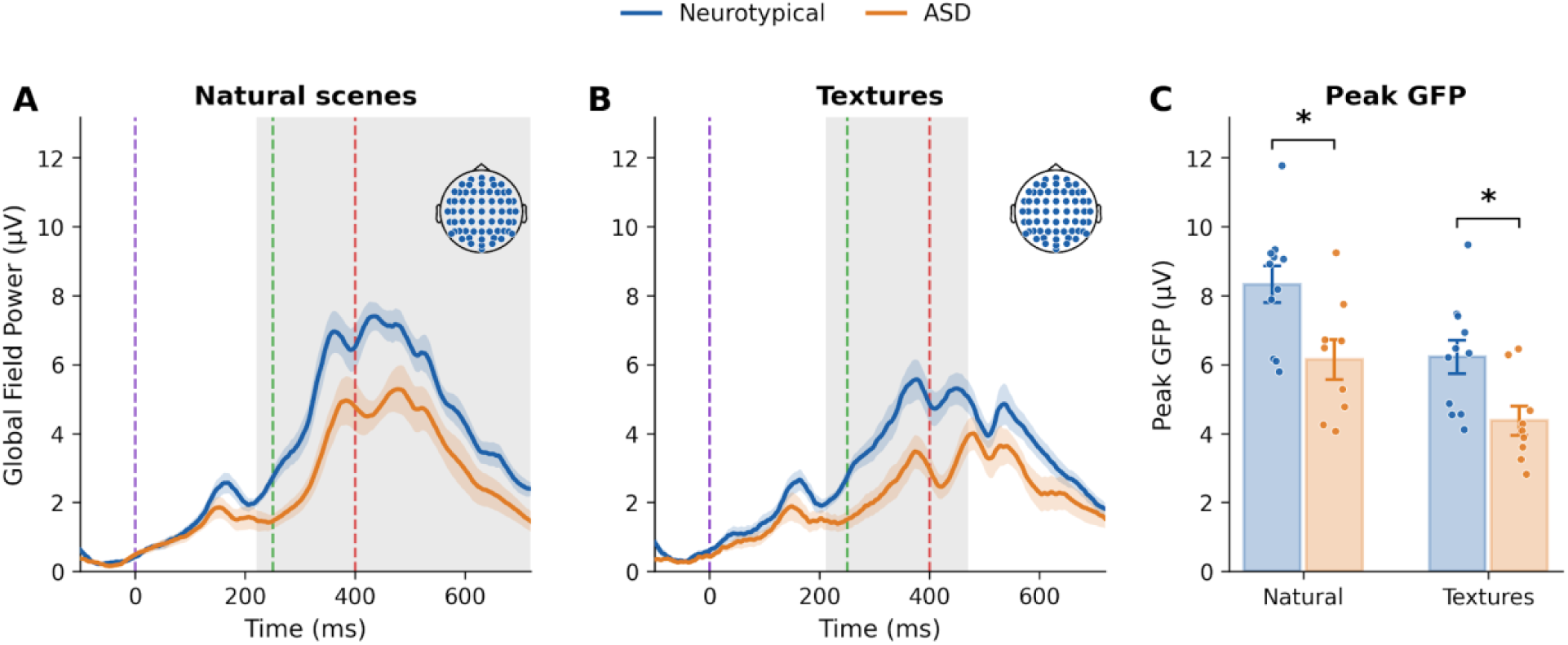
Global field power was higher in neurotypical participants during natural-scene and texture segmentation. (A,B) GFP, defined as the spatial standard deviation across all electrodes, is shown from −100 to 720 ms for natural scenes (A) and textures (B), averaged across participants within each cohort (neurotypical, blue, n = 11; ASD, orange, n = 9); bands denote SEM. Dashed vertical lines mark cue onset (0 ms), image onset (250 ms), and image offset (400 ms); inset heads show the 64-electrode montage. Gray regions mark significant between-cohort clusters from the permutation test across time: 221–719 ms for natural scenes (p = 0.006) and 211–471 ms for textures (p = 0.012). (C) Peak GFP was defined for each participant as the maximum within 0–500 ms. Dots show participants and bars show group mean ± SEM. Peak GFP was higher in neurotypical than ASD participants for natural scenes (8.33 ± 0.53 vs 6.14 ± 0.57 µV; Welch p = 0.012) and textures (6.22 ± 0.49 vs 4.37 ± 0.42 µV; Welch p = 0.010). *p < 0.05.

**Figure 12.**
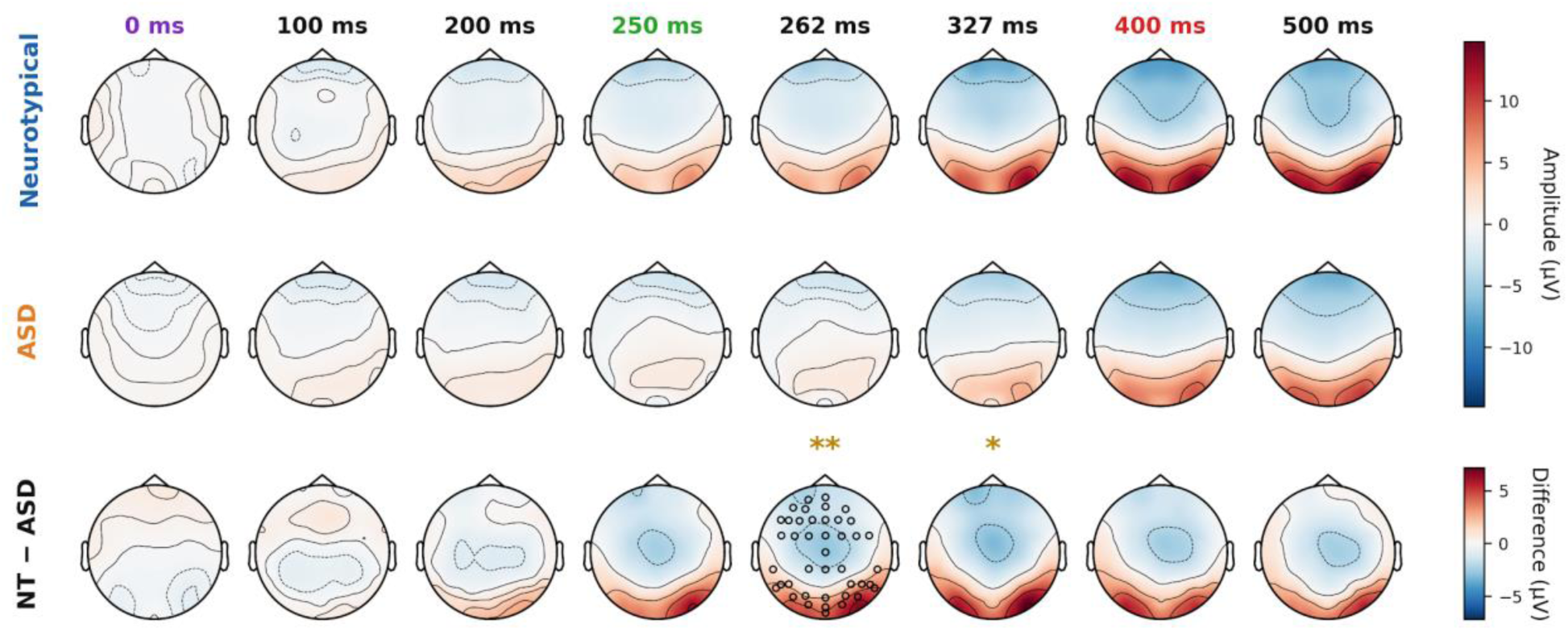
Scalp topography of the cohort difference. Grand-average topographies for Neurotypical (top) and ASD (middle) and their difference (NT − ASD, bottom) at eight latencies. Column-label colors denote trial events, as in Figs. 10–11: cue onset (0 ms, purple), image onset (250 ms, green), image offset (400 ms, red). The amplitude scale (top two rows, ±14.8 µV) and the difference scale (bottom row, ±7.2 µV) are plotted separately; note the smaller range of the difference maps. Asterisks mark within-latency NT − ASD spatial clusters after correction across electrodes: ** 262 ms (p = 0.002) and * 327 ms (p = 0.011); only 262 ms remained significant after Bonferroni correction across the 12 sampled latencies. Outlined electrodes indicate membership in the significant spatial cluster at 262 ms, not significance at individual electrodes.

Because fixation was not enforced, cue-evoked saccades and differences in gaze allocation may contribute to the observed ERP patterns and help explain differences from previous studies. Our concurrent eye-tracking measurements provide an opportunity to assess these contributions through trial-aligned analyses of gaze and fixation-related responses^77^. A further, more speculative possibility is that cohort differences in predictive or integrative processing contribute to these responses. For example, participants may differ in how spatial cues and prior expectations shape anticipatory activity before image onset^17,67,76^. This hypothesis remains to be tested rather than established by the present ERP findings. Future studies combining trial-aligned gaze analyses with control conditions using identical visual events but task-irrelevant cues will help distinguish contributions from looking behavior, general visual processing, and segmentation-related predictions.

Our analysis of ERPs focused on occipital and parieto-occipital channels, because they capture mainly activity in regions canonically involved in early visual encoding, contour integration, and visually driven recurrence^62,78^. This choice was further supported by our topographical results: across analyses, the most reliable cohort differences emerged in O/PO channels (Supplementary Table 1), indicating that these are the spatial locations where cohort-level effects are expressed in this task at this level of analysis. Nonetheless, based on prior studies of segmentation in ASD^14,37^, we also expect differences in a complex neural network that includes higher order cortical regions and in the interaction between brain regions. It is possible that those differences were not apparent in our analysis because we aligned activity to trial onset: the variability of response time across trials could hide effects that are time-locked to the responses.

Our paradigm provides a significant methodological bridge between highly controlled stimuli (illustrated here with the texture-based stimuli) and naturalistic image perception. Previous segmentation work has relied almost exclusively on textures or simplified stimuli^4,13,14,36,37,67,79–81^. To our knowledge, this is the first study to examine visual segmentation using both textures and natural images within the same experimental framework. Including textures is not simply a comparison condition, but rather it allows us to use the fact that texture stimuli have a well-defined ground-truth segmentation, which serves as an internal positive control. The fact that participants in both cohorts recovered these ground truth segments confirms they understood and performed the task correctly, which in turn gives us confidence that the segmentation maps obtained for natural images (where no single ground truth exists) reflect genuine perceptual organization rather than task artifacts. This distinction is critical to test whether cohort differences in segmentation ability generalize across stimulus types and whether the underlying neural mechanisms differ when the perceptual demands increase. This framework also makes it possible to incorporate images with and without social stimuli, and with interactions between people, allowing tests of whether social content distinctly modulates segmentation processes, a question motivated by literature on altered social and face-processing mechanisms in ASD^37,40^.

Importantly, our approach is not limited to natural scenes or basic textures. It can also be applied to classical segmentation stimuli such as Wechler block figures^14,82^, coarse face/house images^37^, or Kanisza figures^36^. A major strength of our paradigm is that it generates full segmentation maps rather than binary global judgments. This capability is critical because these stimuli have been central to decades of work on perceptual grouping and illusory contours, yet prior studies have been limited to inferring perception from categorical same/different or present/not present stimuli^78,83–88^. For example, using our method, theories proposing that individuals with ASD perceive the Kanizsa triangle or other illusory contours differently can be tested directly: in addition to inferring perception from binary behavioral judgements, we can now also measure the full segmentation map and its associated uncertainty along perceptual boundaries. Applying our method to these canonical stimuli would provide a bridge between insights gained from highly controlled artificial stimuli and the more ecologically valid perceptual processes measured in naturalistic scenes.

An additional strength of this paradigm is the high density of observations obtained for each participant, including hundreds of segmentation judgments per trial image. This density supports within-participant analyses of how trial-level behavior, gaze, and neural activity covary, while also characterizing heterogeneity that group averages may obscure. Prior work with the same behavioral paradigm showed that segmentation dynamics tracked participant-specific spatial biases in natural-image segmentation.^89^ Combining these behavioral measurements with concurrent EEG and eye tracking could therefore test how participant-specific perceptual dynamics are expressed in neural and oculomotor signals, including in autism.

An additional perspective is the extent to which the observed neural cohort differences reflect the segmentation task specifically versus general image viewing. Because the current paradigm did not include a passive-viewing or task-irrelevant control condition, the present data cannot disambiguate segmentation-specific activity from responses associated with general image viewing. The ERP and GFP differences preceded the delayed keypress and therefore are unlikely to reflect motor execution, but their timing and successful task performance do not establish a specifically perceptual rather than sensory origin. A future control task using identical visual events with task-irrelevant cue locations will be needed to isolate segmentation-related neural activity.

Our ASD sample consisted of primarily high-functioning individuals. It remains unclear whether similar neural or behavioral patterns would be observed in individuals with differing levels of symptom severity, or, if they would be able to understand and complete the task in its current form. Once a basic characterization of these processes is established in the cohorts considered here, an important next step will be to adapt the paradigm for a broader range of participants, including younger children and individuals with severe intellectual disabilities.

Making the task more developmentally accessible through shorter blocks, simplified instructions, or reducing motor demands, will allow us to test whether the same segmentation mechanisms generalize across ability levels and developmental stages. Such adaptations would also create opportunities for longitudinal studies that track the development of perceptual segmentation within the same participants over time.

## ACKNOWLEDGMENTS

We thank Theo Vanneau for guidance on developing the EEG preprocessing pipeline and Tringa Lecaj and Dennis Cregin for helping with recruitment and data collection. R.C.C. was supported by the National Institutes of Health (EY031166); the Rose. F. Kennedy Intellectual and Developmental Disabilities Research Center. R.C.C. and S.M. were supported by the Simons Foundation International (SFI-AN-AR-Pilot-00009855). Support for recruitment and phenotyping of participants was provided by the Human Clinical Phenotyping Core of the NICHD funded Rose. F. Kennedy Intellectual and Developmental Disabilities Research Center (P50 HD105352).

## Author contributions

D.K.A.W. contributed to collecting the data; preprocessing and analyzing the EEG data, analyzing the choice and reaction time data; drafting and writing the manuscript.

T.B. contributed to preprocessing and analyzing the choice and reaction time data; writing the manuscript.

C.B. contributed to preprocessing and analyzing the gaze data and implemented the Information Gain analysis; writing the manuscript

S.M. contributed to acquiring funding; designing the experiments; implementing the study design; writing the manuscript.

R.C.C. contributed to conceiving the project; acquiring funding; designing the experiments; implementing the study design; collecting the data; preprocessing and analyzing the choice and reaction time data; drafting and writing the manuscript.

All authors approved the final manuscript for submission.

## SUPPLEMENTARY INFORMATION

**Supplementary Figure 1.**
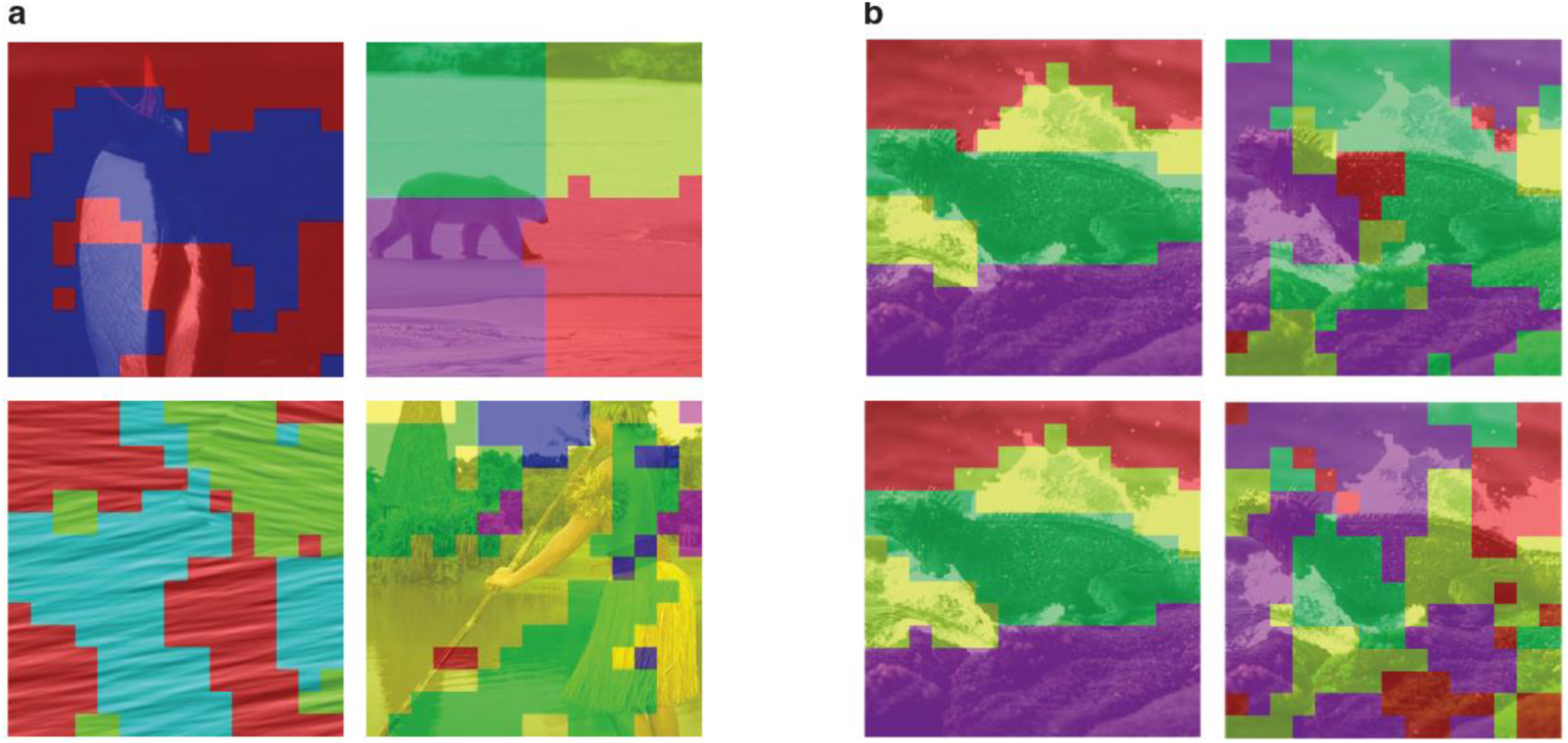
(**a**) Examples of maps from blocks excluded from the analysis. We debriefed participants after each block and asked them to indicate verbally the segments they perceived in the image. By comparing those descriptions with the maps we reconstructed numerically, we identified experimental blocks in which the participants did not perform the same-different task: in those cases, the maps do not correspond to the description nor to any meaningful segments. In addition, in one block (top right) the instructions were misinterpreted: on debriefing the participant explained that, because there was no right or wrong answer, they decided to segment the image in four quadrants rather than based on how they perceived it. This is evident from the reconstruction. (**b**) For two participants (rows), examples of maps from blocks included in the analysis (left) and as a control the maps reconstructed after randomly shuffling the order of the participants’ responses across trials (right). Trial shuffling destroys the relation between the spatial cues in a trial and the corresponding response from the participant. The resulting maps lack spatial structure and appear unrelated to the image content or to the verbal description. This control analysis supports that the reconstructed maps from the original data are meaningful and not artifactual.

**Supplementary Figure 2.**
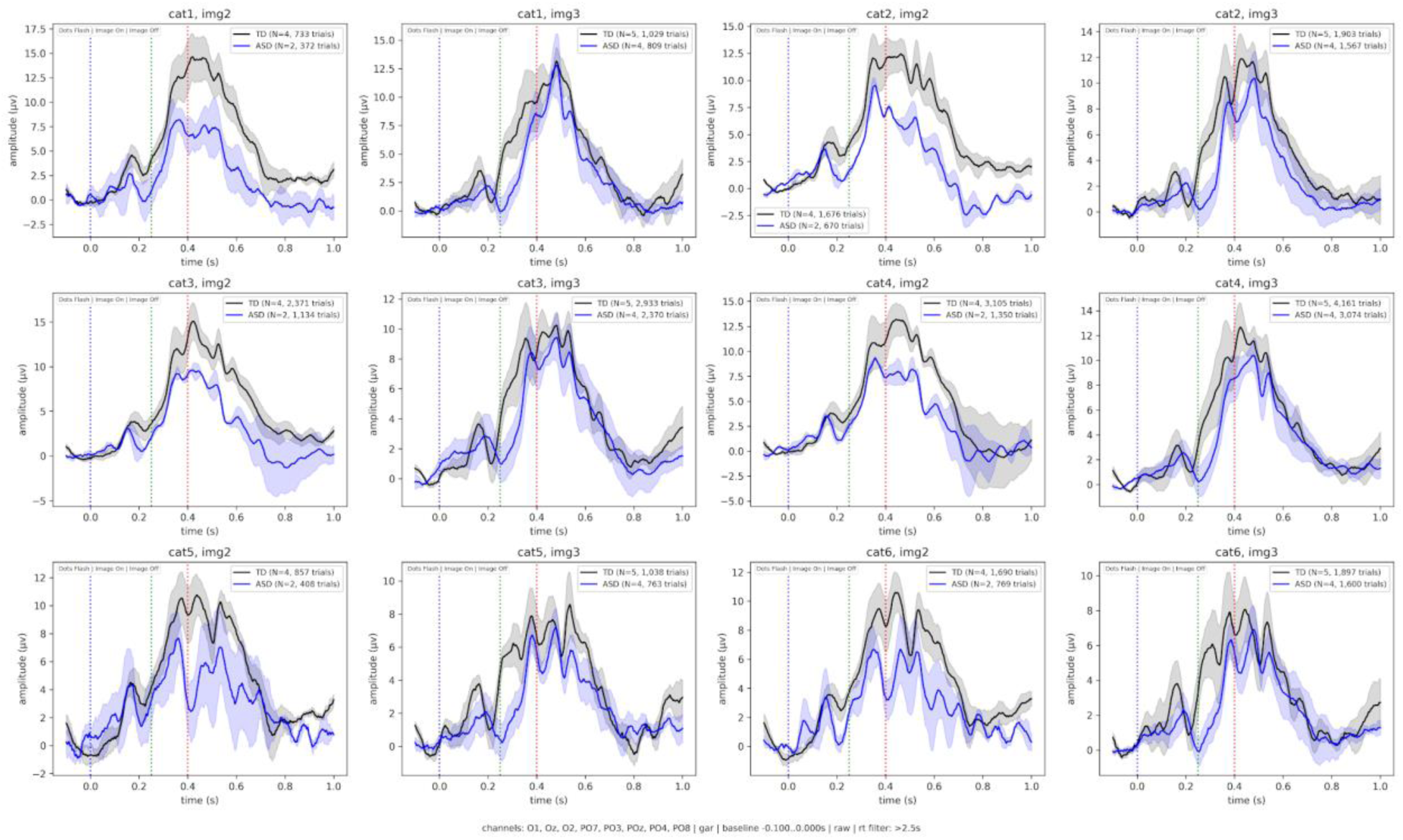
Event-related potentials (ERPs) comparing neurotypical (NT; black) and ASD (blue) groups across all experimental images. Grand-average waveforms over occipital and parieto-occipital channels (O1, O2, Oz, PO7, PO3, POz, PO4, PO8) are shown descriptively for each category-image combination (categories 1–6, images 2–3). Vertical dashed lines mark cue onset (0 ms, blue), image onset (250 ms, green), and image offset (400 ms, red).

**Supplementary Table 1.** Statistical comparison of NT vs ASD groups across time windows: Global Field Power (GFP), Region of Interest (ROI), and pointwise electrode analyses for Grand Average and Category 3/Image 3 conditions.

| Grand Average |  |  |  |  |  |  |
| --- | --- | --- | --- | --- | --- | --- |
| GFP p-value | Significance | ROI p-value | ROI Effect Size (d) | N Significant Electrodes | ROI Electrodes (p-values) | Top 5 Other Electrodes (p-values) |
| 0.001347848 | ** | 0.001374147 | 2.073323491 | 29 | PO8 (0.0005), O2 (0.0005), Oz (0.0022), O1 (0.0024), PO4 (0.0025), PO7 (0.0130) | TP8 (0.0001), FC3 (0.0001), P10 (0.0001), Iz (0.0001), P8 (0.0002) |
| 0.000927539 | *** | 0.002253886 | 1.953481221 | 36 | PO8 (0.0004), O2 (0.0012), PO4 (0.0029), O1 (0.0055), Oz (0.0058), PO7 (0.0090), POz (0.0489) | P10 (0.0000), P8 (0.0002), TP8 (0.0002), FC3 (0.0003), Iz (0.0003) |
| GFP p-value | Significance | ROI p-value | ROI Effect Size (d) | N Significant Electrodes | ROI Electrodes (p-values) | Top 5 Other Electrodes (p-values) |
| 0.001190091 | ** | 0.007212254 | 1.661282998 | 36 | PO8 (0.0012), O2 (0.0075), PO7 (0.0079), PO4 (0.0089), O1 (0.0113) | P10 (0.0001), P8 (0.0008), Iz (0.0008), TP8 (0.0010), FC3 (0.0012) |
| 0.010600359 | * | 0.078357955 | 1.008740636 | 16 | PO8 (0.0040), PO7 (0.0328) | P10 (0.0010), TP8 (0.0021), P8 (0.0023), P6 (0.0073), FC1 (0.0086) |
| 0.002969443 | ** | 0.006106457 | 1.689522889 | 22 | PO8 (0.0002), O2 (0.0006), Oz (0.0058), O1 (0.0107), PO4 (0.0242), PO7 (0.0266) | P10 (0.0000), TP8 (0.0003), P8 (0.0004), Iz (0.0005), FC1 (0.0014) |
| Single Image (Cat3/Img3) |  |  |  |  |  |  |
| GFP p-value | Significance | ROI p-value | ROI Effect Size (d) | N Significant Electrodes | ROI Electrodes (p-values) | Top 5 Other Electrodes (p-values) |
| 0.020045715 | * | 0.016229924 | 2.158762807 | 18 | PO8 (0.0045), PO4 (0.0054), O2 (0.0089), Oz (0.0243), O1 (0.0310), POz (0.0324) | P6 (0.0009), P10 (0.0011), P8 (0.0011), TP8 (0.0012), P4 (0.0019) |
| 0.00590006 | ** | 0.016391845 | 2.111154807 | 20 | PO8 (0.0031), PO4 (0.0046), O2 (0.0083), POz (0.0345), Oz (0.0360), O1 (0.0465) | P10 (0.0002), P8 (0.0003), P6 (0.0003), P4 (0.0015), TP8 (0.0018) |
| 0.026273168 | * | 0.053596504 | 1.572244959 | 11 | PO8 (0.0115), O2 (0.0116), Oz (0.0160) | Iz (0.0218), FC3 (0.0251), C2 (0.0261), P8 (0.0280), P10 (0.0290) |

## Footnotes

✝ *<u>Terminology note:</u> There is ongoing discussion within the autism community regarding whether person-first language (“individuals with autism”) or identity-first language (“autistic individuals”) is preferred. Current preferences vary across autistic adults, self-advocates, clinicians, and researchers. In this paper, we tend to use person-first language, consistent with conventions in clinical and neuroscience research, while recognizing and respecting that many in the autism community prefer identity-first language. Our choice is not intended to imply a value judgment, and we acknowledge the validity of both approaches*.

